# Genome-wide profiling identifies the genetic dependencies of cell death following EGFR inhibition

**DOI:** 10.1101/2025.04.04.647273

**Authors:** Sydney A. Porto, Gavin A. Birdsall, Nicholas W. Harper, Megan E. Honeywell, Michael J. Lee

## Abstract

EGFR is a proto-oncogene that is mutationally activated in a variety of cancers. Small molecule inhibitors targeting EGFR can be effective in slowing the progression of disease, and in some settings these drugs even cause dramatic tumor regression. However, responses to EGFR inhibitors are rarely durable, and the mechanisms contributing to response variation remain unclear. In particular, several distinct mechanisms have been proposed for how EGFR inhibition activates cell death, and a consensus has yet to emerge. In this study, we use functional genomics with specialized analyses to infer how genetic perturbations effect the drug-induced death rate. Our data clarify that inhibition of PI3K signaling drives the lethality of EGFR inhibition. Inhibition of other pathways downstream of EGFR, including the RAS-MAPK pathway, promote growth suppression, but not the lethal effects of EGFR inhibitors. Taken together, our study reveals the first “reference map” for the genome-wide genetic dependencies of lethality for EGFR inhibitors.

## INTRODUCTION

Molecularly targeted therapies have revolutionized cancer treatment. Targeted therapies can distinguish between normal cells and cancer cells, ideally leading to highly selective killing of cancer cells with low toxicity. These treatments are typically designed to inactivate oncogenes in growth factor signaling pathways. While these treatments have been successful in many settings, responses are generally not durable (1). Previous research on the failures of targeted therapies has focused on understanding mechanisms of drug resistance (2). However, less attention has been applied to determining the mechanisms of drug sensitivity, and in particular, why inhibition of a growth factor oncogene promotes tumor cell death.

Some of the earliest and most well-studied molecularly targeted therapies are epidermal growth factor receptor (EGFR) tyrosine kinase inhibitors (TKIs). First-generation EGFR inhibitors, including gefitinib and erlotinib, reversibly bind to EGFR and inhibit the binding of ATP, consequently abolishing the receptor’s tyrosine-kinase activity (3). These two drugs had remarkable success in patients with EGFR-mutant non-small cell lung cancer (NSCLC) (4–6), and erlotinib was ultimately approved as a first-line treatment for NSCLC patients harboring EGFR exon 19 deletion or exon 21 (L858R) substitution mutations (7). However, the success of first-generation EGFR inhibitors was limited due to the emergence of drug-resistant populations, and in particular, the appearance of the T790M ‘gatekeeper’ mutation (8–10). Second and third-generation EGFR inhibitors were developed, in part, to address this issue of resistance, although novel resistance mutations have continued to arise (11). Even though EGFR inhibitors have been clinically relevant and effective for decades, and research has focused extensively on mechanisms of resistance to EGFR inhibition, there remains a need to further understand why these targeted treatments work.

In cases where EGFR-TKIs are effective treatment strategies, it remains unclear how inhibiting EGFR activates cell death. Several major signaling pathways downstream of EGFR control cell proliferation and survival, most notably the RAS-RAF-MEK-ERK and PI3K-AKT-mTOR pathways (12). Previous studies have suggested that EGFR inhibition might cause cell death through either the RAS pathway, the PI3K pathway, a combination of both, or a separate downstream pathway (13–15). Given this ambiguity, there is a need to better define how EGFR-TKIs cause cell death. A detailed understanding of the mechanisms of lethality for EGFR TKIs may aid in identification of patients who are likely to respond to these drugs, and may help to predict novel resistance mechanisms or more effective drug combinations.

Functional genomics can be used to identify the genetic dependencies of drug sensitivity, and theoretically, also the mechanisms of drug-induced cell death. Advances in CRISPR-Cas9-mediated genome editing have enabled high-throughput, quantitative characterization of all genes and/or gene regulatory elements (16). When conducted in the context of a drug treatment, these data are often referred to as a “chemo-genetic profile” (17). Chemo-genetic profiling has been effective at characterizing the genetic dependencies for “cell fitness”. However, these chemo-genetic profiles are not consistently able identify mechanisms of drug-induced lethality. The central issue is that growth rate variation among clones confounds the interpretation of chemo-genetic profiling data, often masking the degree to which genetic perturbations alter cell death. To address this issue, we recently developed a **<u>M</u>**ethod for **<u>E</u>**valuating **<u>D</u>**eath **<u>U</u>**sing a **<u>S</u>**imulation-assisted **<u>A</u>**pproach (**MEDUSA**) (18). MEDUSA employs computational simulations that model clonal dynamics to reveal the combination of growth and death rates that created an observed drug response, and how these rates are altered by each single gene knockout.

In this study, we examine the response of EGFR-mutant NSCLC cells to the first-generation EGFR-TKI erlotinib and the third-generation inhibitor osimertinib. We determined that, although these drugs consistently induce growth inhibition, these inhibitors do not consistently activate cell death, even among NSCLC cell lines harboring the same EGFR mutations. Drug-induced changes in gene expression could not account for the observed differences in lethality across genetic backgrounds. Using chemo-genetic profiling with a MEDUSA-based analysis, we uncovered the genetic determinants of cell death following EGFR inhibition. MEDUSA highlighted that the lethality of EGFR inhibitors depends on inhibiting the PI3K-AKT-mTOR signaling branch. Importantly, the insights generated by MEDUSA were distinct from the insights generated using conventional chemo-genetic profile analyses. Taken together, these data provide a complete reference for the genetic dependencies of EGFR inhibitor-induced death.

## RESULTS

### EGFR inhibitors rarely activate cell death while consistently promoting growth inhibition in EGFR-mutant NSCLC

EGFR signals through several downstream pathways that regulate cell survival and proliferation. Four canonical pathways downstream of EGFR are: the RAS-RAF-MEK-ERK, PI3K-AKT-mTOR, PLCγ, and JAK-STAT pathways (Figure 1A) (12, 19). Based on literature precedent, inhibition of any one of these pathways could independently activate cell death. Indeed, prior studies have specifically connected EGFR inhibitor-driven cell death to the inhibition of either one or multiple pathways, and a consensus model has yet to emerge. For instance, several studies have suggested that EGFR inhibitor-induced cell death is driven through inhibition of the RAS-RAF-MEK-ERK pathway, due to loss of ERK-mediated BIM phosphorylation (20–22). BIM is a pro-apoptotic protein whose degradation is regulated by ERK, such that upon ERK inhibition, BIM protein accumulates and may activate cell death (13). In contrast, other studies suggest that the combination of inhibiting both the RAS-RAF-MEK-ERK and PI3K-AKT-mTOR pathways drives apoptosis, with inhibition of the former leading to the upregulation of BIM, and inhibition of the latter leading to the downregulation of the anti-apoptotic protein Mcl-1 (14). Other pathways downstream of EGFR have also been connected to cell death. For instance, nuclear import of PKCδ has been shown to activate apoptosis (23, 24). Furthermore, STAT1 and STAT3 have both pro- and anti-apoptotic functions (25, 26). Given these numerous and conflicting mechanisms describing how pathways downstream of EGFR contribute to cell death activation, the mechanisms of drug sensitivity following EGFR inhibition remain unclear.

**Figure 1.**
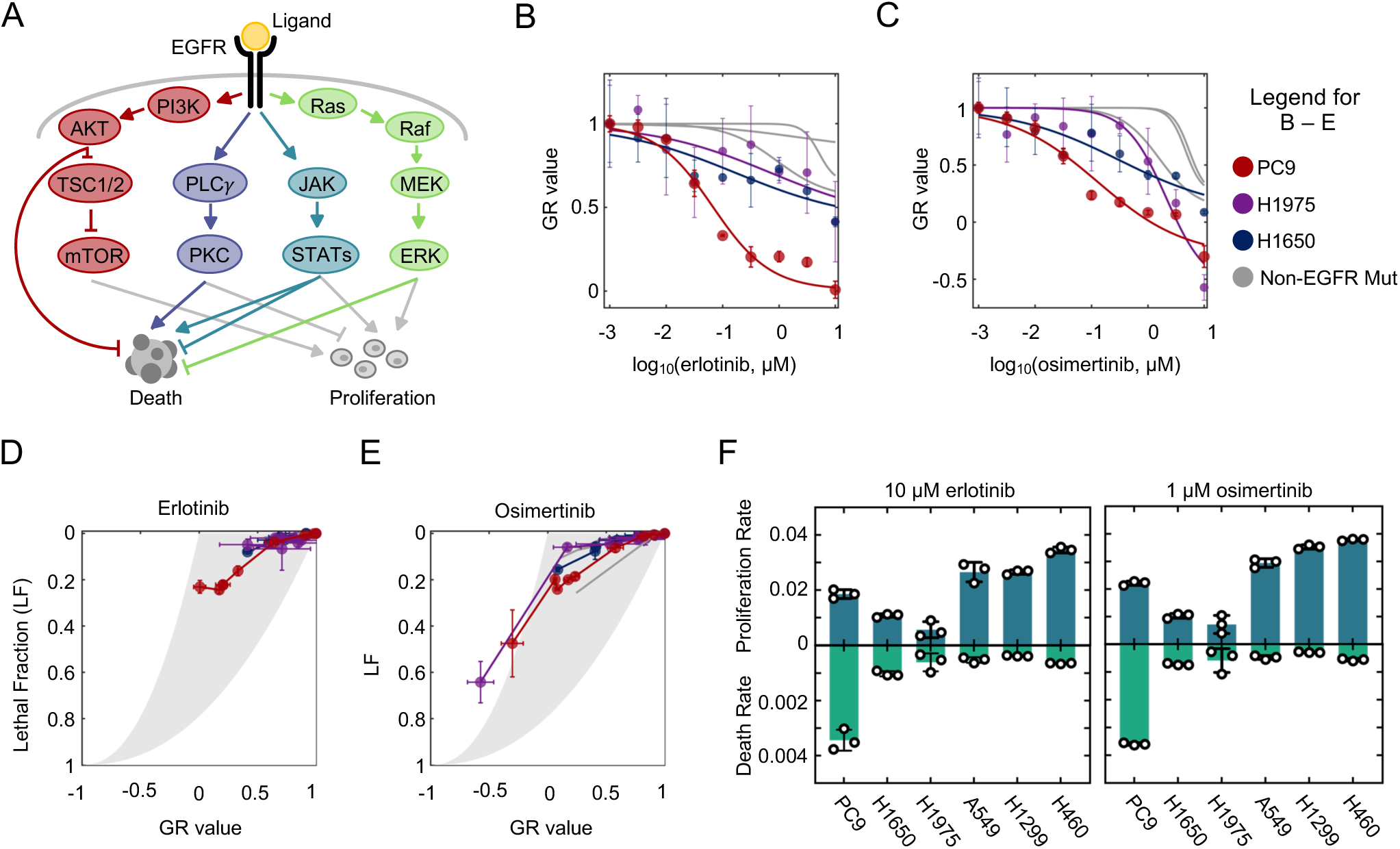
EGFR inhibitors rarely activate cell death, even in EGFR-mutant NSCLC cells. (**A**) Simplified schematic of key signaling pathways that control survival and proliferation downstream of EGFR. (**B**) Erlotinib sensitivity of PC9, H1975, H1650, and three non-EGFR mutant NSCLC cells, evaluated 72 hours following erlotinib exposure. GR value reports the net population growth/shrinkage rate, relative to untreated cells. (**C**) As in (B) but following exposure to osimertinib. (**D**) Drug GRADE analysis for following 72-hour exposure to erlotinib. Cell lines are colored as in panel (B) and (C). GRADE analysis juxtaposes the GR value with the drug-induced Lethal Fraction (LF) to visualize the drug-induced coordination between growth and death. Each cell line was exposed to erlotinib at an 8-point dose titration, as in panels (B) and (C). (**E**) As in panel (D) but following exposure to osimertinib. (**F**) Proliferation rate (doublings/hour) and death rate (LF/hour) for cell lines treated with 10 µM erlotinib or 1 µM osimertinib. Values calculated from GRADE analysis. For all panels with error bars, data are the mean ± S.D. for *n =* 3 independent biological replicates.

To evaluate mechanisms of lethality following EGFR inhibition, we began by profiling EGFR inhibitor sensitivity in a panel of EGFR-mutant and EGFR wild-type NSCLC cell lines. We focused on the first-generation inhibitor, erlotinib, and the third-generation inhibitor, osimertinib. Because we tested these drugs across cells with varied growth rates, drug sensitivities were evaluated using the normalized growth rate inhibition (GR) value (27). GR values account for growth rate variation between cells, therefore allowing for more accurate comparisons of drug responses across cell lines. GR values are scaled between 1 and −1 where a GR value between 0 and 1 report an expanding population, a value of 0 represents complete cytostasis, and a value between 0 and −1 indicates a shrinking population. Previous studies have generally suggested that EGFR wild-type NSCLCs are less sensitive to EGFR inhibitors than NSCLCs containing activating mutations in EGFR (28). Consistent with these prior results, we find that the cell line with the highest sensitivity to erlotinib and osimertinib was the EGFR mutant cell line PC9 (EGFR *delE746-A750*) (Figure 1B-C). Unexpectedly, however, other EGFR mutant NSCLC cells, H1650 (*delE746-A570*) and H1975 (L858R, T790M), were only modestly more sensitive than the EGFR wild-type NSCLCs that we profiled (Figure 1B-C). These results were consistent with prior studies reporting that PC9 cells are particularly sensitive to EGFR inhibition (29).

We next sought to further characterize the increased EGFR inhibitor sensitivity in PC9 cells. The GR value reports the net population growth rate at each dose of drug; however, GR values alone cannot clarify the relative contributions of growth inhibition versus cell death activation to the observed drug response. To determine the proportional contributions of growth inhibition and cell death activation following exposure to EGFR inhibitors, we used the drug GRADE analysis approach (Figure 1D-E) (30). The GRADE method juxtaposes the GR value and the drug-induced lethal fraction (LF), which enables a visualization of how growth inhibition and death activation are coordinated by a given drug. GRADE reveals that PC9 cells respond to erlotinib and osimertinib using a “high GRADE” response (Figure 1D-E). This feature indicates that a large fraction of the observed response to EGFR inhibition results from activation of cell death. In contrast, the other cell lines – including all other EGFR mutant NSCLCs – have low GRADE responses to EGFR inhibition, revealing that these cells respond to EGFR inhibitors only with slowed proliferation rates, and without activation of cell death. A notable exception is observed for high doses of osimertinib, which appear to kill all cells, likely due to off-target effects at these concentrations (Figure 1E).

To more precisely quantify how growth arrest and cell death contributed to the observed drug responses, we used the GRADE analysis to infer the quantitative combinations of growth rates and death rates that created the observed drug responses for each tested cell type. For PC9 cells, quantitative inference from GRADE reveals that erlotinib and osimertinib reduce the proliferation rate to 0.019 population doublings per hour, slightly less than half of the untreated growth rate of PC9; these drugs also activated death with an average rate of 0.34% per hour (Figure 1F). In all other contexts, EGFR inhibition reduced the growth rate of these cells to only ∼ 60-90% of the untreated growth rate, without causing significant levels of drug-induced death (Figure 1F). Therefore, EGFR inhibitors cause variable responses across the cell lines included in this panel, even among the three EGFR-mutant NSCLCs tested. With respect to the six cells used in this study, erlotinib and osimertinib appear to be uniquely lethal in PC9 cells.

### Differences in drug-induced changes in gene expression do not account for variable lethality across genetic backgrounds

In the panel of cells that we profiled, elevated levels of lethality following EGFR inhibition were only observed in PC9 cells, so we next aimed to further characterize why EGFR inhibition was more effective in promoting cell death in this setting. To address this, we examined EGFR inhibitor-induced death in PC9 and H1650, two cell-lines harboring the same EGFR mutation (*delE746-A750*), which have varied death rates following EGFR inhibition (Figure 1F). Kinetic analysis of cell death revealed that high dose erlotinib (10 µM) kills ∼30-40% of PC9 cells over a 72-hour window (Figure 2A). Osimertinib was significantly more potent, with a 100-fold lower dose resulting in a similar level of lethality (Figure 2A). To explore the subtype of cell death activated by erlotinib or osimertinib in PC9 cells, we quantified the drug response in the presence and absence of z-VAD, an inhibitor of apoptotic caspases. For both erlotinib and osimertinib, z-VAD significantly inhibited EGFR inhibitor-induced lethality, and inhibitors of other pathways did not alter the level or the timing of cell death (Figure 2B). These data suggest that EGFR inhibition activates an apoptotic cell death, which is consistent with prior studies (31–33). Importantly, kinetic evaluation in H1650 cells revealed that the modest death rate that is observed is not drug-induced, but rather results from a high-level of background cell death in these cells (Figure 2C). Thus, PC9 experiences drug-induced lethality when exposed to EGFR inhibition, while H1650 does not.

**Figure 2.**
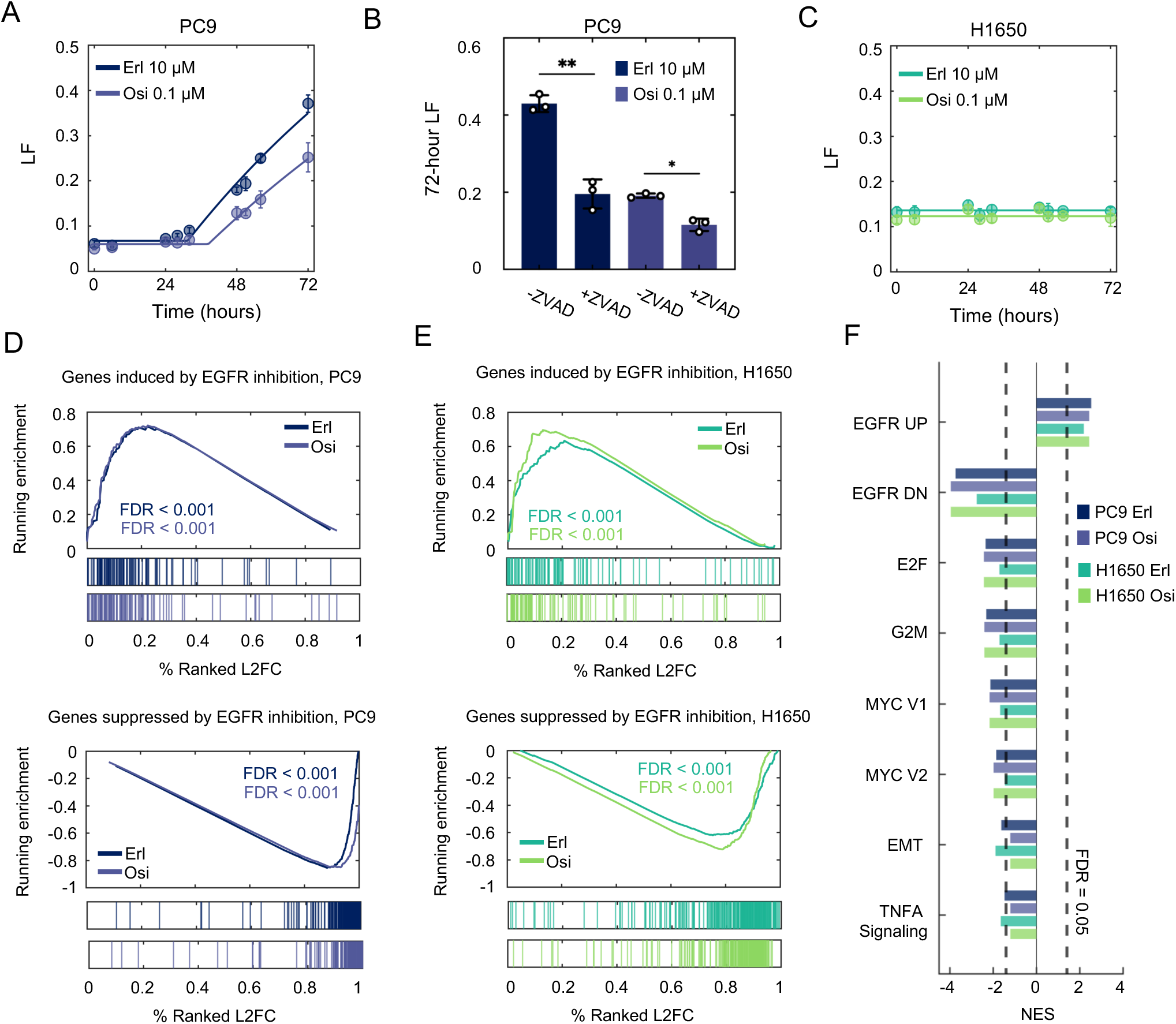
Drug-induced changes in gene expression cannot explain the observed variation in the lethality of EGFR inhibitors. (**A**) Cell death kinetics in PC9 cells, measured using the FLICK assay following exposure to either 10 µM erlotinib or 0.1 µM osimertinib. (**B**) 72-hour lethal fraction for PC9 cells exposure to 10 µM erlotinib or 0.1 µM osimertinib, in the presence or absence of 50 µM z-VAD. \**p* < 0.01 and \*\**p* < 0.001 using a two-sided *t*-test. (**C**) As in panel (A) with but in H1650 cells. (**D**) Gene set enrichment analysis (GSEA) for erlotinib- and osimertinib-induced gene expression changes in PC9 cells. Gene signatures used are from msigDB (“Kobayashi EGFR Signaling 24-hour UP” and “Kobayashi EGFR Signaling 24-hour DN”). FDR-adjusted p-values are based on 1000 permutations of gene sets. (E) As in panel (D) but for H1650 cells. (**F**) GSEA for drug-treated conditions compared to untreated samples for PC9 and H1650. Significantly changed enriched Hallmark Genesets shown. For all panels with error bars, data are the mean ± S.D. for *n =* 3 independent biological replicates.

Given that PC9 and H1650 cells have such different responses to EGFR inhibition, we next aimed to determine if the lethality observed in PC9 – or the lack of lethality in H1650 – could be explained by differences in drug-induced gene expression changes. To test this, we performed RNA-sequencing in PC9 and H1650 cells 36 hours after exposure to erlotinib or osimertinib (the time of death onset in PC9 cells). We then performed gene set enrichment analysis (GSEA) to quantify and visualize the drug-induced expression profiles (34). Erlotinib- and osimertinib-induced changes in gene expression are well characterized; thus, we compared our results to previously annotated genetic signatures for the transcriptional effects of EGFR inhibition in EGFR-mutant NSCLC cells in different contexts. For instance, Kobayashi et al. previously characterized gene expression signatures comprised of 102 or 253 genes that were up- or down-regulated, respectively, specifically in NSCLC cells that are sensitive to EGFR inhibition, but not in insensitive cells (35). In PC9 cells exposed to either erlotinib or osimertinib, the transcriptional effects that we observed were similar to those observed previously in EGFR-mutant NSCLCs that are sensitive to EGFR inhibiton (Figure 2D). Surprisingly, however, nearly identical gene expression changes were also observed following erlotinib and osimertinib treatment in H1650 cells, despite the relative lack of sensitivity and complete lack of drug-induced cell death that we observe following EGFR inhibition (Figure 2E).

To explore the similarities and differences between the transcriptional responses in PC9 and H1650 more broadly, we also identified other gene signatures that were impacted by EGFR inhibition in these cells. Again, for both cell lines, and for both EGFR inhibitors, similar transcriptional responses were observed across all hallmark gene signatures, and overall, we failed to identify any differences in the transcriptional responses to EGFR inhibition that could clarify the observed differences in drug-induced lethality (Figure 2F). Taken together, these data show that in EGFR inhibitor-sensitive PC9 cells and EGFR inhibitor-insensitive H1650 cells, EGFR inhibition alters gene expression in a similar manner, despite the stark differences in drug-induced lethality. Thus, drug-induced gene expression changes appear not to account for the differences in lethality following EGFR inhibition.

### MEDUSA-based analysis reveals mechanism of lethality following EGFR inhibition

Since changes in gene expression were not able to explain mechanisms of EGFR-inhibitor driven death in PC9 cells, we sought to clarify the mechanisms of lethality by identifying the genetic dependencies of EGFR inhibition-induced cell death in PC9 cells. We performed a genome-wide single gene knockout screen using the TKOv3 library (Figure 3A) (36). Cells were treated with either 10 µM erlotinib or 0.1 µM osimertinib for 72 hours, which resulted in ∼30-40% of the population dying (Figure 2A). To examine the quality of our chemo-genetic profiling data, we first explored the degree of correlation between our biological replicates (37). These data revealed a high correlation between replicates, both at the level of sequencing read counts and at the level of drug-induced fold change, suggesting a high degree of biological reproducibility (Counts: r = 0.97; Fold change: r = 0.82). Furthermore, when comparing the “T0” input samples to the untreated samples at the end of the assay, we observed core essential genes dropping out over time, suggesting that our chemo-genetic profiling data could accurately recover expected genetic dependencies.

**Figure 3.**
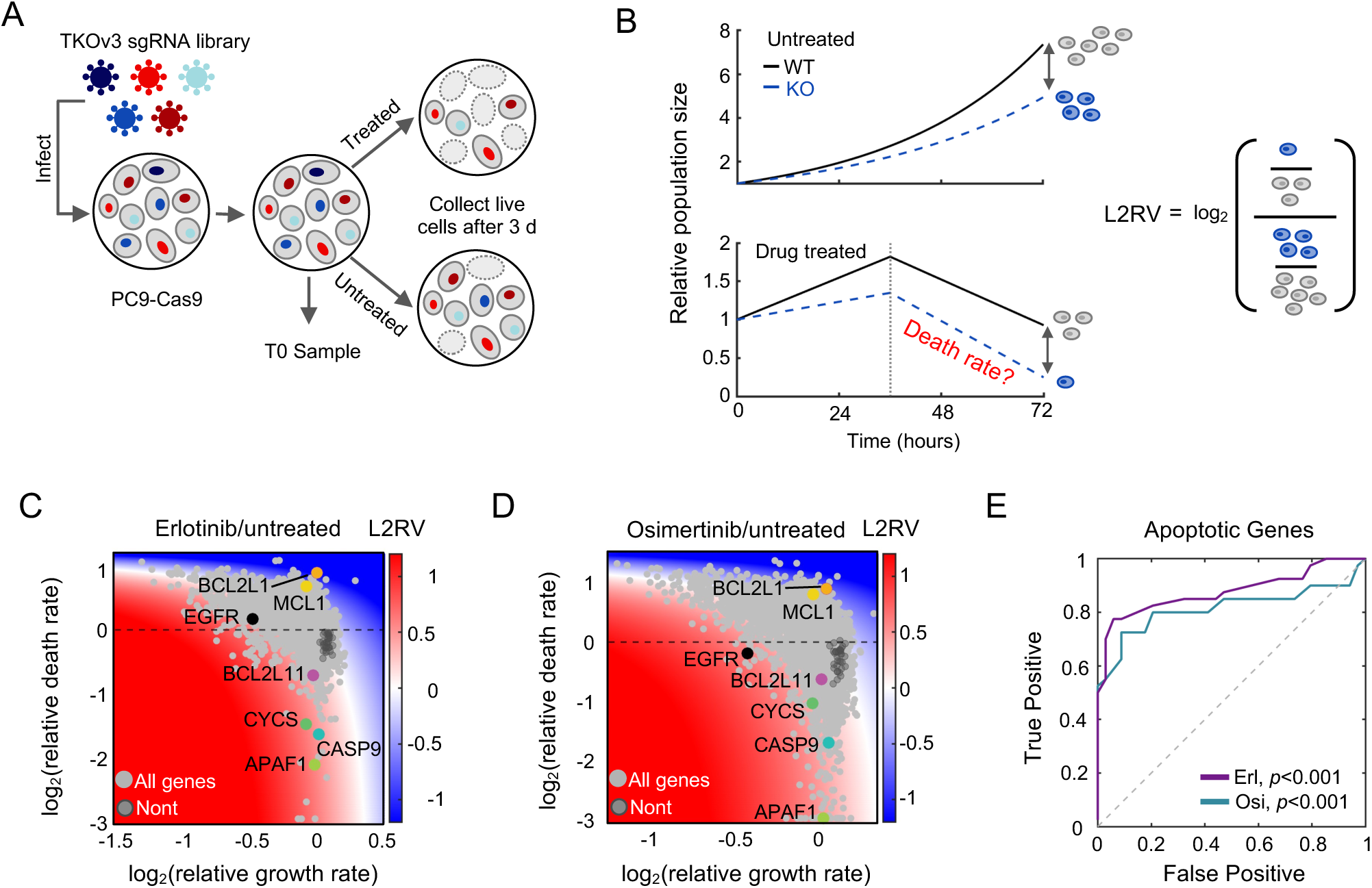
Chemo-genetic profiling with MEDUSA-based analysis reveals the genetic dependencies of lethality following EGFR inhibition. (**A**) Schematic of chemo-genetic profiling using a pooled sgRNA library. (**B**) Model showing the relationship between drug-induced growth and death rates, and the conventional chemo-genetic profiling analysis (log2-Relative Viability, L2RV). (**C**) Gene-level chemo-genetic profiling with MEDUSA analysis for PC9 cells treated with 10 µM erlotinib for 72 hours. Non-targeting genes are shown in dark gray. Several known apoptotic regulators and EGFR are highlighted. (**D**) As in panel (C) but following 0.1 µM osimertinib exposure. (**E**) ROC analysis for MEDUSA-inferred death rates of known apoptotic genes. Empiric *P* value based on AUC and bootstrapping with 1,000 iterations.

To identify genes that regulate EGFR inhibitor-induced cell death, we employed the **<u>M</u>**ethod for **<u>E</u>**valuating **<u>D</u>**eath **<u>U</u>**sing a **<u>S</u>**imulation-assisted **<u>A</u>**pproach (**MEDUSA**) (18). Conventional chemo-genetic profiles rely on examining the relative abundance of each clone, comparing treated and untreated populations (Figure 3B).

These methods – which are effectively comparing four dynamic populations, each with their own growth and death rates – fail to capture how genetic perturbations affect drug-induced cell death, due to the confounding effects of variable proliferation rates and death rates within the other populations (18). The MEDUSA method solves this issue using comprehensive simulations of the drug-treated and untreated populations growing and dying at varied rates, to map the observed fold change data to a single pair of growth and death rates.

The outcome of a MEDUSA-based analysis is a computational inference of the drug-induced growth and death rates for each knockout clone in a chemo-genetic profiling dataset. To evaluate the accuracy of our analysis, we began by determining MEDUSA’s effectiveness at identifying genes involved in intrinsic apoptosis, as apoptotic regulation is reasonably well-studied. For instance, knocking out any component of the apoptosome – which is comprised of caspase-9 (*CASP9*), cytochrome c (*CYCS*), and APAF-1 (*APAF1*) – should compromise the drug-induced death rate (38). As expected, our MEDUSA-based analysis revealed knocking out *CASP9*, *CYCS*, or *APAF1* significantly rescued erlotinib or osimertinib induced death rates (Figure 3C-D). Furthermore, these genetic dependencies would not have been clearly identified using a conventional “log-fold change” based analysis. MEDUSA also identified potent negative regulators of apoptosis, such as Bcl-xL (*BCL2L1*) and Mcl-1 (*MCL1*), which, when knocked out, sensitized cells to the lethal effects of both erlotinib and osimertinib (Figure 3C-D). Consistent with prior literature which has highlighted the central importance of the pro-apoptotic protein BIM (*BCL2L11*) (20–22), BIM knockout suppressed erlotinib-induced cell death, although the effect was somewhat modest (Figure 3C). BIM knockout did not, however, significantly decrease drug-induced lethality in the context of osimertinib (Figure 3D). Overall, MEDUSA-based analysis was effective at recovering known apoptotic regulatory genes (Figure 3E).

Quantitatively, our chemo-genetic profiling data for erlotinib- or osimertinib-induced death were strongly correlated, suggesting an overall similarity between the mechanisms of lethality for these two drugs (r = 0.71). In total, our analysis identified 676 genes that significantly modulate death rate in response to erlotinib treatment, and 674 genes that modulate death in response to osimertinib. 359 of these apoptotic regulatory genes were shared as regulators of both drugs. Taken together, these data represent the first consensus map of the genetic dependencies of lethality following EGFR inhibition in EGFR-mutant PC9 cells.

### Inhibition of PI3K signaling drives lethality following EGFR inhibition

Having determined that our screen was successful in identifying apoptotic regulatory genes, we next sought to use these data identify which signaling pathways were contributing to death following EGFR inhibition. We began by mapping the genes that regulate lethality of EGFR inhibitors onto the canonical signaling pathways downstream of EGFR (Figure 4A). We saw that knocking out several components of the PI3K pathway altered the erlotinib-induced death rate. Indeed, knocking out activators in the pathway, including PDK1 and ILK, sensitized PC9 cells to death (Figure 4A). Likewise, knocking out inhibitors of the PI3K pathway, including PTEN and both members of the TSC complex, rescued EGFR inhibitor-induced cell death (Figure 4A).

**Figure 4.**
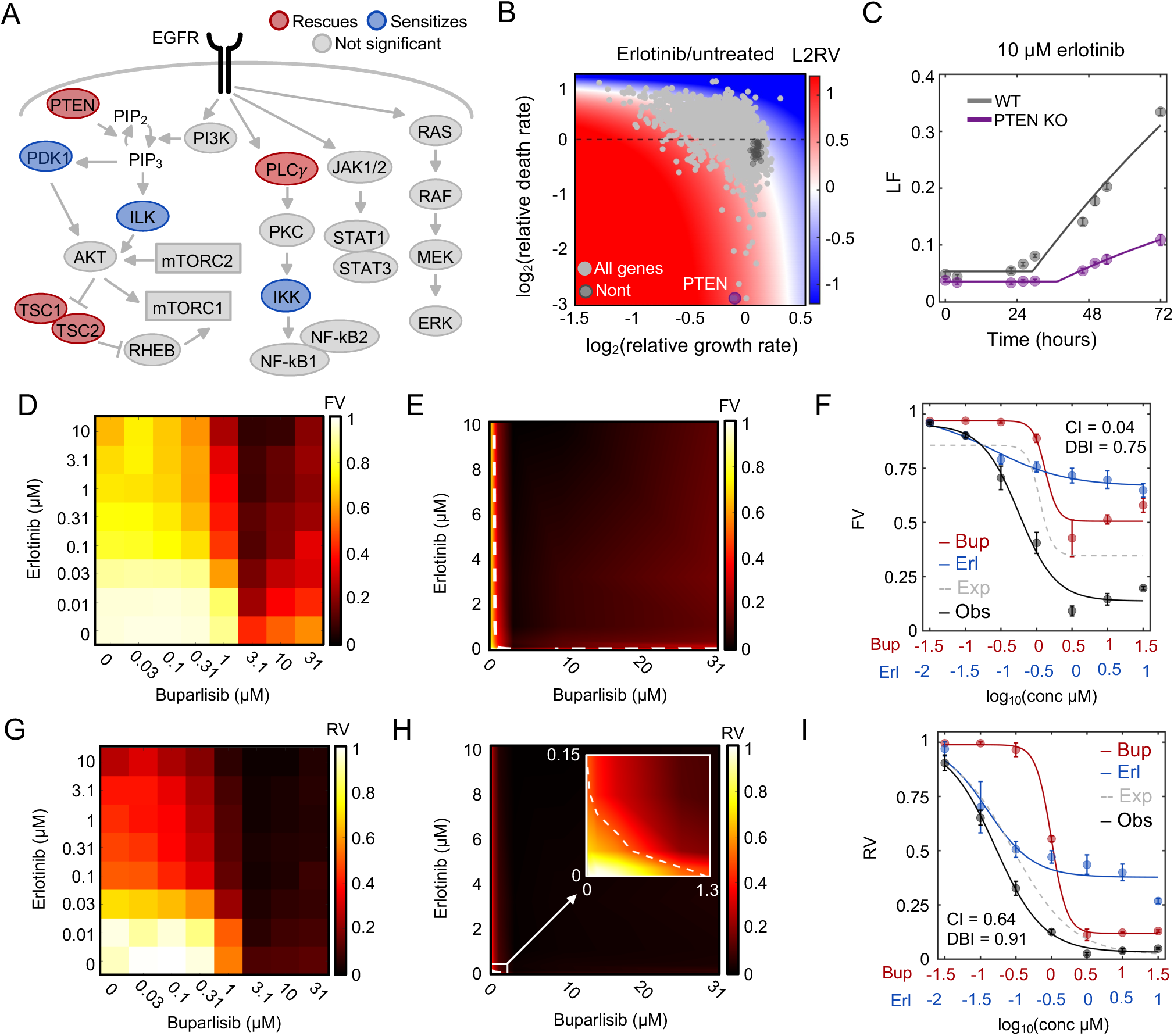
Inhibition of PI3K signaling drives the lethality of EGFR inhibition. (**A**) Schematic of signaling pathways downstream of EGFR. Death regulatory genes identified in MEDUSA are highlighted. Genes whose knockout sensitizes death are shown in blue, while genes whose knockout rescues death are shown in red. (**B**) Gene-level chemo-genetic profile for PC9 cells treated with erlotinib. *PTEN*, the top hit based on the MEDUSA death rate, is highlighted. (**C**) Validation of MEDUSA-inferred death rate for *PTEN*. Erlotinib-induced death evaluated using the FLICK assay. (**D**) Dose titration of erlotinib and buparlisib in PC9 cells. Heatmap is scaled by the mean fractional viability of 3 biological replicates following 72 hours of drug treatment. (**E**) Isobologram analysis for the data in (D). The dashed line represents the erlotinib and buparlisib combinations that result in 50% response (50% isobol). (**F**) Dose curve for erlotinib and buparlisib at fixed ratio dosing. The expected dose curve in the case of additivity is shown in addition to the observed combination. The Chou-Talalay Combination Index (CI) and the Deviation from Bliss Independence (DBI) are shown. (**G**-**I**) As in (D-F), but data are the conventional relative viability metric, rather than a death-specific metric. For all panels with error bars, data are the mean ± S.D. for *n =* 3 independent biological replicates.

Among the canonical signaling pathways downstream of EGFR, death regulation appears to be unique to the PI3K pathway. We did not see the same abundance of significant hits in the other signaling pathways downstream of EGFR. Genes within the PLCγ, JAK/STAT, and RAS pathways generally did not contribute to EGFR inhibition-induced cell death, and overall, these pathways were not significantly enriched for apoptotic regulatory genes. It is also worth noting, however, that genetic redundancy, particularly within the RAS pathway, may lead to a gene within these pathways failing to score in a screen focused on single gene knockouts (39, 40).

To validate the functional significance of the PI3K pathway for promoting EGFR-induced cell death, we focused initially on genetically validating single gene knockouts identified by our MEDUSA-based analysis as regulators of the lethality of erlotinib and osimertinib. PTEN is a phosphatidylinositol-3,4,5-trisphosphate (PIP_3_) phosphatase which is a potent negative regulator of PI3K signaling (41, 42). Our MEDUSA-based analysis revealed that *PTEN* knockout strongly suppressed EGFR inhibitor-induced lethality (Figure 4B). To validate this insight, we knocked out *PTEN* using targeted sgRNAs, and these data demonstrate that knocking out *PTEN* decreased the erlotinib-induced cell death to approximately one-third of the wild-type death rate (Figure 4C).

To further validate the importance of PI3K signaling for promoting the lethality of EGFR inhibitors – we next sought to quantify the extent to which chemically inhibiting PI3K signaling alters erlotinib-induced lethality. To address this question, we tested erlotinib in the presence and absence of the PI3K inhibitor, buparlisib, with each drug evaluated across a large dose titration, at all pairwise dosing combinations (Figure 4D). We scored single drug and combinatorial drug responses using the “fractional viability” metric (FV, live cells divide by the total of live and dead cells). FV differs from conventional response metrics, which compare treated- and untreated populations, and importantly, FV is a death-specific measurement (43). Using this death-specific analysis, we found that erlotinib and buparlisib induce moderate lethality as single agents, but that the lethality is exacerbated when these drugs are used in combination (Figure 4D).

To characterize the interaction between erlotinib and buparlisib in PC9 cells, we used well-validated measures of drug-drug interactions (i.e., drug “synergy” or “antagonism”) (43, 44). We began by visually inspected the combinatorial response data using the Loewe isobologram convention. In the Loewe model, one would expect the lines connecting similarly efficacious doses to produce linear “isobols” if the two drugs integrate additively. However, combinations of erlotinib and buparlisib produced overwhelmingly concave isobols, indicative of a strong drug synergy (Figure 4E). To quantify the degree of drug synergy, we used the Chou-Talalay Combination Index (CI), which revealed a CI of 0.04, equivalent to a 25-fold reduction of the dose required for 50% lethality when compared to a dose additivity model (Figure 4F). Similarly strong synergy was observed when these data were evaluated using the Bliss Independence reference model (DBI, Figure 4F).

To further validate our MEDUSA-based analysis, we next focused on whether inhibition of PI3K signaling was specifically promoting the lethality – and not the growth suppression – of EGFR inhibitors. To address this question, we re-analyzed our erlotinib-buparlisib drug combination data using the conventional “relative viability” (RV) metric (Figure 4G). RV is based on the size of the drug-treated population, compared to an untreated control population. In contrast to FV, which directly score drug-induced death, RV is sensitive to changes in proliferation, but relatively insensitive to changes in cell death (45, 46). Quantitative evaluation of these data revealed a modest, and statistically insignificant, trend towards drug synergy, with the erlotinib-buparlisib combination resulting in a 50% reduction in RV at a dose within 2-fold of the expectation based on dose additivity (CI = 0.64, Figure 4H-I). Taken together, these data confirm our MEDUSA-based inferences and highlight that inhibition of the PI3K signaling pathway drives the lethality of EGFR inhibition, whereas the other signaling pathways downstream of EGFR appear to primarily regulate proliferation.

### Genetic dependencies of lethality for EGFR inhibition are distinct from the genetic dependencies of cell fitness or cell proliferation

Given that EGFR inhibitors are well studied, we next determined whether the insights generated by our MEDUSA-based inference of death regulation are distinct from insights generated using other methods. For instance, many studies have explored mechanisms regulating EGFR inhibitor efficacy by studying EGFR inhibitor-induced changes in gene expression (35, 47, 48). We identified more than 1,400 genes that are differentially expressed following exposure to erlotinib; however, we observed no relationship between the differentially expressed genes and the ∼600 genes that regulate erlotinib-induced death (Figure 5A). Similarly, expression changes were not correlated with the growth regulatory function of each gene (Figure 5B). These observations mirror those made previously in the context of yeast stress responses, which also highlight that genes induced by a stress tend to be distinct from the genes that are functionally required for the observed response (49). Collectively, these observations demonstrate that the insights generated by gene expression-based analysis of EGFR inhibition are unrelated to the genetic dependencies of drug-induced lethality.

**Figure 5.**
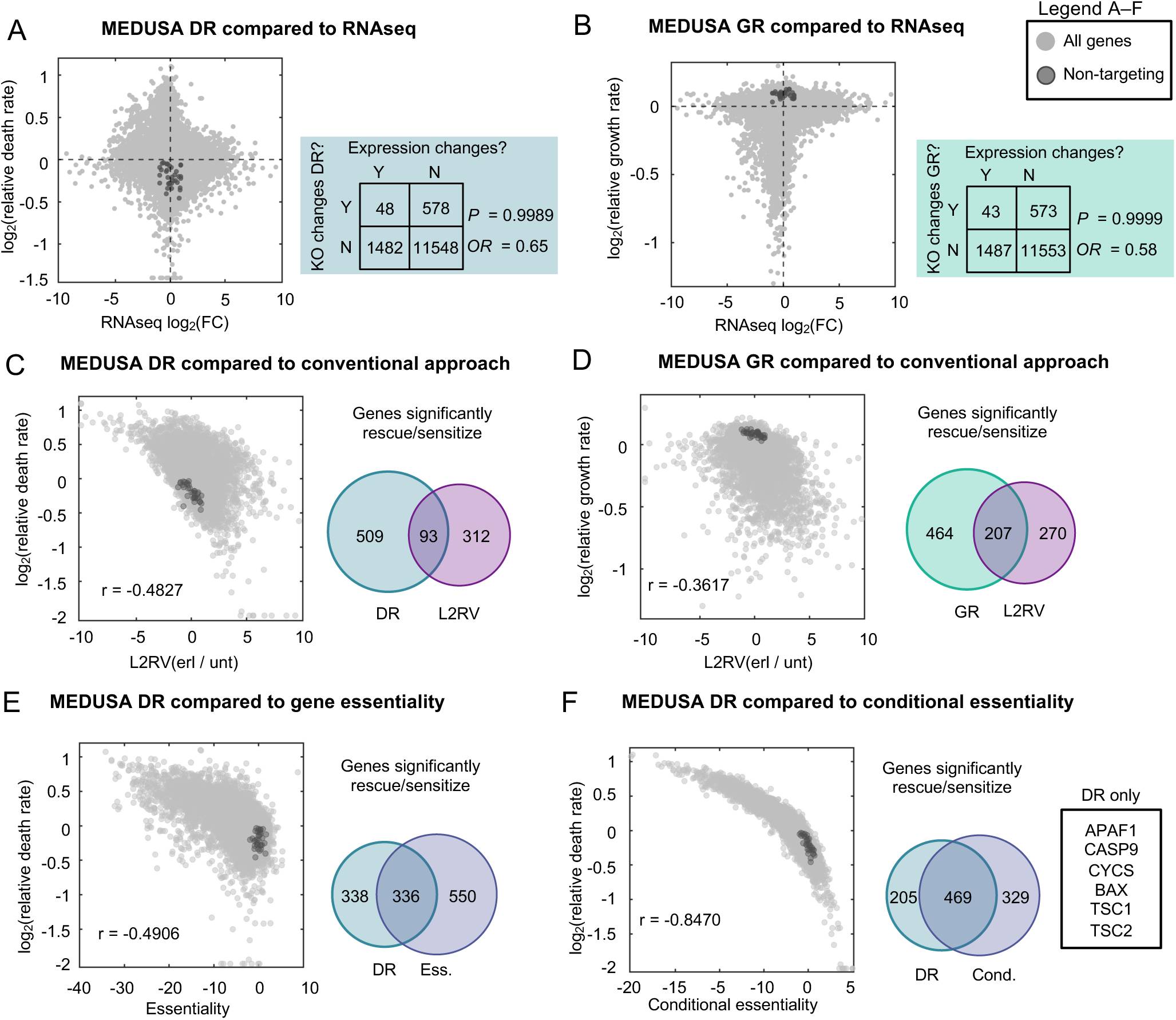
Genetic dependencies of lethality are distinct from the genetic dependencies of cell fitness or cell proliferation. (**A**) Comparison of MEDUSA-inferred death rate and drug-induced gene expression changes. Shaded regions of the graph genes that are significantly differentially expressed and whose deletion significantly alters the drug-induced death rate. Odds Ratio (OR) and p-value based on Fisher’s Exact Test to determine the relationship between gene expression changes and death regulatory function. (**B**) As in panel (A) but for the MEDUSA-inferred growth rate compared to differential gene expression. **(C)** Comparison of MEDUSA death rate to conventional chemo-genetic profiling analysis (L2RV). Venn diagrams show the relative number and relationship between “hits” recovered by each analysis strategy. (**D**) As in panel (C) but for MEDUSA GR. (**E**-**F**) Further comparison of the MEDUSA DR metric to other conventional metrics used in functional genomics and/or chemo-genetic profiling. (E) Comparison between MEDUSA DR and gene essentiality. (F) Comparison between MEDUSA DR and conditional essentiality. Several genes only recovered by the MEDUSA analysis, but not conditional essentiality, are highlighted in (F).

In addition to the numerous studies that have used gene expression changes to infer mechanisms of drug action for EGFR inhibitors, several other studies have explored EGFR inhibitors using chemo-genetic profiling (50–52). Previous chemo-genetic profiling studies, however, have invariantly used analysis methods that fail to accurately identify death regulatory genes (generally a log-fold change-based analysis, comparing the relative abundance of each knockout clone in drug treated versus untreated conditions). Indeed, no prior studies were effective at enriching for apoptotic genes, even though PC9 cells activate apoptotic death when exposed to EGFR inhibitors. We therefore compared the MEDUSA-inferred relative death rates (log_2_ death rate, L2DR) to the genetic dependencies based on a conventional log-fold change analysis (log_2_ relative viability, L2RV). We observe a weak negative correlation of r = −0.48 between the L2DR and L2RV metrics (Figure 5C).

Importantly, the MEDUSA-inferred growth rates for drug treated cells were also not well-correlated with conventional a L2RV-based analysis; however, the MEDUSA-inferred growth rates were strongly correlated with measures of gene essentiality (e.g., growth fitness when measuring endpoint versus T0 input samples, r = 0.976) (Figure 5D). The strong positive correlation between MEDUSA-inferred growth rates and gene essentiality scores further demonstrates the accuracy of MEDUSA-based growth and death rate inferences, and suggests that gene essentiality is generally a measure of proliferative fitness, not death regulation.

Although L2RV or similar methods are the most common methods used in chemo-genetic profiling studies, gene essentiality (i.e., L2RV when comparing untreated and T0 samples) and/or gene conditional essentiality, (i.e., L2RV when comparing drug treated and T0 samples) can also be used to assess genetic dependencies (36). Similar to what was observed for conventional L2RV-based analysis of treated vs. untreated conditions, we observed a negative correlation between the MEDUSA-inferred death rate and gene essentiality score, suggesting that essential genes tend to increase the erlotinib-induced death rate when knocked out (r = −0.49, Figure 5E). However, despite the correlation between these measures, gene essentiality was not an effective strategy for mechanisms of drug-induced cell death, as the essentiality data was not effective at identifying an apoptotic mechanism of action for erlotinib.

Similar data were observed when comparing MEDUSA-inferred death rates and the conditional gene essentiality, although the correlation between these two measures was very strong (r = −0.85, Figure 5F). This is not surprising, given that conditional essentiality is effectively scoring the relative fitness in a drugged condition for each gene knockout compared to the wild-type cells, which is a more biologically interpretable comparison than the conventional analysis. Nonetheless, conditional essentially also fails to replicate the sensitivity and accuracy of MEDUSA-inferred death rates. For instance, MEDUSA alone identifies key regulators of apoptosis, including all subunits of apoptosome: APAF1, CASP9, and CYCS (Figure 5F). Overall, as was true for all other analytical strategies, conditional essentiality fails to clearly identify an apoptotic mechanism of action for erlotinib. Taken together, these data demonstrate that MEDUSA-inferred growth and death rates provide quantitative insights that are not captured using other conventional analysis methods.

## DISCUSSION

In this study, we explored the mechanisms by which EGFR inhibitors activate cell death in the context of EGFR-mutant NSCLC cells. Using death-specific assays – including MEDUSA, to identify death regulatory genes – our data highlight the central importance of inhibiting the PI3K signaling pathway. Inhibition of other signaling branches, including the RAS-MAPK pathway inhibited the proliferation of PC9 cells, without promoting high levels of cell death. Our data and analyses reveal that these phenotypes are challenging to interpret using conventional methods, including RNAseq, which has been widely used to infer mechanisms of drug function. Thus, the data and analytical strategies presented here represent a valuable resource for understanding how EGFR inhibitors activate cell death, and how to potentiate these responses.

More generally, from the perspective of chemo-genetic profiling, our data demonstrate that conventional chemo-genetic profiling analysis methods are neither measures of drug-induced death or drug-induced growth inhibition, but rather an amalgam of these traits across four populations (i.e., knockout and wild-type, in treated and untreated conditions). Notably, similar analytical strategies are used effectively in the context of gene essentiality profiling (53). A critical difference is that the relative fold change analysis used in gene essentiality profiling compares the dynamics of the knockout versus wild-type population, compare to a static “T0” population, which is used only to normalize for differences in starting population size. When applied in the context of chemo-genetic profiling, rather than gene essentiality, this analytical strategy confounds the interpretation, both from the perspective of death activation versus growth inhibiting drug effects, and regarding the objective response. For instance, a positive L2FC can be generated from a knockout population that is resistant to a drug, or from a knockout population that grows slowly in the absence of drug, but does not alter the drug response.

One unexpected finding from this study is that EGFR inhibitors do not typically activate high levels of cell death, even in the context of EGFR-mutant NSCLCs. Indeed, among the EGFR-mutant cell lines that are widely used, we identified only a single example, PC9 cells, which respond to EGFR inhibition with robust activation of apoptotic cell death. At least three plausible interpretations exist. First, because our study was performed using common tissue culture conditions, micro-environmental influences may change the probability and levels of drug-induced cell death. Second, it remains possible that the clinically observed responses to EGFR inhibitors do not result from cell autonomous forms of lethality, such as cell intrinsic apoptosis. The methods of data collection and analysis that we used can accurately report all cell autonomous types of apoptotic and non-apoptotic cell death, but immune cell-mediated forms of cell death were not profiled. Finally, a third interpretation could be that EGFR inhibitor-induced lethality is indeed uncommon. PC9 cells may represent a “super responder” type clone. Future studies should aim to distinguish between these, and potentially other, possibilities.

Nonetheless, our data demonstrate that high levels of drug-induced cell death are possible following EGFR inhibition. The genetic dependencies that we identify should help to clarify the sources of response variation and should reveal rational strategies to improve the efficacy and durability of EGFR inhibitors.

## Acknowledgements

Funding for this project was provided by the National Cancer Institute (NCI) grant U01CA265709 (to MJL). The authors report no conflicts of interest.

## METHODS

### Cells and reagents

PC-9 (NSCLC) cells were a gift from J. Pritchard (Penn St.). NCI-H460 cells were acquired from the Green Laboratory (UMass Chan Medical School). A549, H1650, H1975, and H1299 cells were obtained from the American Type Culture Collection (ATCC). Cells were maintained at a low passage number, below 20 from the original vial. A549 cells were grown in DMEM (Corning, 10-017-CV), and all other cells were grown in RPMI 1640 medium (Corning, 10-040-CV). Each media was supplemented with 10% FBS (Peak Serum, PS-FB2), and penicillin–streptomycin (Corning, 30-002-CI).

SYTOX Green (S7020) was purchased from Thermo Fisher Scientific. Erlotinib hydrochloride salt was purchased from LC Laboratories (catalog no. E-4007), osimertinib was purchased from MedChemExpress (catalog no. HY-15772), and buparlisib was purchased from MedChemExpress (catalog no. HY-70063). Z-VAD-FMK was purchased from ApexBio (catalog no. A1902).

### FLICK-based analysis of drug response

The FLICK assay was conducted as described previously (54, 55). Briefly, cells were seeded at a density of 2,000 cells per well in 96-well plates (Greiner Bio-One, 655090) in 80-90 µL of media and incubated overnight. Cells were drugged with their respective doses of drug or vehicle in 2-5 µM of SYTOX Green media (Thermo Fisher Scientific, S7020) such that the final volume of each well was 100 µL. Dead cell fluorescence was measured with a Tecan Spark (ex: 503, em: 524), and gain was selected for each cell line as previously described (55). A T0 plate was lysed at the start of each assay by adding 1.5% Triton X-100 (Thermo Fisher Scientific, BP151-100) in PBS and incubating at 37°C for 3 hours. At the end of the experiment, all plates were lysed using 1.5% Triton X-100 in PBS and incubated as before. The permeabilized plates provided the total number of cells at the start and end of the assay. From the total cell fluorescence and dead cell fluorescence, the live cell fluorescence can be inferred. This information allowed us to calculate RV, FV, LF, and GR (27). A custom MATLAB script was used to generate dose-response curves and LF kinetic curves as described previously (54).

### Gene expression analysis with RNAseq

To analyze drug-induced changes in gene expression, PC9 and H1650 cells were seeded at 200,000 cells per well in a six-well plate and incubated overnight. Cells were then treated with either 0.1% DMSO or 10 µM erlotinib. T0 samples were collected at the time of drugging. After 36 hours, the remaining samples were washed with 1 mL of PBS and removed from the plate with 500 µL of 1.5% trypsin. The cells were then spun down at 500 xg for 5 minutes, washed in PBS, and spun down again under the same conditions. The pellets were snap frozen in liquid nitrogen and stored at –80°C. RNA was extracted according to the manufacturer’s instructions in the Qiagen RNeasy Mini Kit. Each sample had a concentration of at least 180 ng/µL with 260/280 ratios of 2.01-2.05 and 260/230 ratio of 1-1.9. The experiment was completed in duplicate. RNA was sent to Novogene for sequencing. Sequencing results were processed using the UMass Medical School Biocore’s DolphinNext pipeline. DESeq2 in R computed log_2_(fold change) from the generated counts table (treated vs. T0) and computed FDR-adjusted *P* values.

### Generation of PC9-Cas9 cell line

PC9-Cas9 cells were generated using virus containing lenti-Cas9-Blast (Addgene, 52962). Viral infection was performed by plating 2 million PC9 cells per well of a 12-well dish and adding virus and 8ug/ml polybrene (Millipore, TR1003G). Cells were spun at 800xg for 2 hours at 37°C, after which viral media was removed and fresh media was added for cells to recover overnight. The following day, cells were replated and allowed to recover further. 24 hours later, cells were selected using 5 µg/mL blasticidin (ThermoFischer, R21001) for 5 days to generate a stable population. Cas9 expression was verified with a western blot and activity was evaluated using the Broad Institute’s protocol assaying Cas9 Activity with an EGFP reporter assay.

### Chemo-genetic profiling of response to EGFR inhibition

A whole-genome CRISPR screen was performed using the TKOv3 two-vector system (36). PC9-Cas9 cells were transfected with the TKOv3 pooled library via spinfection. For each treatment condition (erlotinib, osimertinib, and DMSO) 45×10^6^ PC9-Cas9 cells were infected, divided into 12-well plates. Each well contained 1.5×10^6^ cells, 15 µL of virus, and 0.8 µL/mL polybrene (Millipore, TR1003G) in a total volume of 2 mL. The plates were centrifuged at 37°C and 830 xg for two hours. Following centrifugation, the media was replaced, and the cells were allowed to recover overnight. The cells were replated on 15 cm plates, incubated overnight, treated with 1 µg/mL puromycin (Corning, 61-385-RA) for 3 days, and then cells allowed to recover 2 days. 270×10^6^ cells were plated in total, with each experimental condition and T0 control having two replicates. On day 0, the treated conditions were drugged with 10 µM erlotinib and the untreated conditions received 0.01% DMSO. The untreated conditions were passaged on day 1. On day 3, live cells from the treated and untreated conditions were collected and frozen.

DNA extraction, PCR and sequencing was performed as described previously (36). In brief, genomic DNA was isolated using the Wizard Genomic DNA purification kit (Promega, A1120) at 10x scale. sgRNA sequences were extracted from each genome by PCR (forward: GAGGGCCTATTTCCCATGATTC, reverse: CAAACCCAGGGCTGCCTTGGAA). A second PCR reaction was used to attach barcodes used for multiplexing. Products from the second PCR reaction were gel-extracted (Qiagen, 28704). Following library balancing, samples were sequenced on the Illumina NextSeq2000.

### Conventional analysis of chemo-genetic profiles (L2RV)

Read quality was verified using FastQC, and non-variable regions were removed to isolate the guide sequences using the FASTX trimmer function. Reads were mapped to the TKOv3 library using Bowtie2 and a single mismatch was allowed. Counts tables were generated and the bottom 5% of guides by base mean were removed from analysis. The L2RV was calculated at the guide level using DESeq2 in R. Guide-level scores were collapsed to gene-level by taking the median. The 142 non-targeting guides were randomly assigned to a ‘gene’ in groups of 4. Gene-level fold changes were z-scored based on the distribution of L2RV scores for the non-targeting genes. An empiric *P* value was determined for each gene via 10,000 iterations, and this score was FDR corrected using the Benjamini–Hochberg procedure.

### MEDUSA analysis

MEDUSA analysis for screen data was performed as previously described (18). Briefly, growth and death rates for PC9 cells in the presence and absence of drug were determined using the FLICK assay and used to model population dynamics. With this model, we simulated all the ways a gene knockout might affect growth and drug-induced death rates. Using these simulated growth and death rates, the relative size of the treated and untreated populations. The conventional L2RV metric was then calculated. The drug-induced growth and death rate for each single-gene knockout was inferred from the observed L2RV, simulations, and relative growth rate of the knockout in the absence of the drug. The MEDUSA-inferred guide level scores were collapsed to the gene-level by taking the mean, and then they were z-scored to the distribution of non-targeting genes. Empiric *P* values were determined via bootstrapping with 10,000 iterations and were FDR corrected.

### Screen validation

Selected guides were cloned into the pX330-puro plasmid using the digestion-ligation protocol from the Zhang laboratory (available on the Zhang laboratory addgene page). PC9 cells were plated at 300,000 cells per well in 6 well dishes in complete medium. The following day, cells were transfected with 1.5 μg of sgRNA-puro constructs using the FuGENE HD Transfection Reagent (Promega, E2311), following the manufacturer’s instructions. The following day cells were replated on 10cm dishes. The next day cells were treated with 1 μg/mL puromycin (Corning, 61-385-RA) and selected for 3 days, washed twice with PBS, and replaced with fresh media and recovery for 2 days. For single cell cloning, selected cells were plated in 96 wells, grown until confluent, expanded into 10 cm dishes, after which, genetic perturbations were validated.

### Data Analysis and Statistics

Unless otherwise noted, data analysis was performed in MATLAB (version R2024a) using built-in functions. Bar graphs with individual data points were generated using GraphPad Prism (version 10.4.1). GSEA analysis was performed using the GSEA 4.3.2 package. Pair-wise statistical comparisons were generated using a two-tailed two-sample t-test. For conventional L2RV and MEDUSA analyses, *P* values were generated by bootstrapping with 10,000 iterations and FDR corrected with the Benjamini-Hochberg procedure.

